# Open-Source, High-Speed and High-Resolution Data Acquisition Platform for Biopotential Recordings and Neural EIT applications

**DOI:** 10.64898/2026.08.06.743202

**Authors:** Enrico Ravagli, Alistair McEwan, Kirill Aristovich

**Author notes:** Author to whom any correspondence should be addressed. **E-mail:**.

## Abstract

**Objective:** Biopotential measurement devices, such as EEG, ECG, and EMG recorders, are available in low-cost, open-source implementations with standard specifications. However, high-end systems remain expensive and predominantly proprietary, limiting accessibility and customization by research laboratories. In addition, neurophysiology techniques such as bioimpedance-based Fast Neural Electrical Impedance Tomography (FN-EIT) also rely on these systems for data acquisition. This work aimed to develop an open-source biopotential recording system using off-the-shelf components that achieves performance comparable to high-end devices.

**Approach:** We designed our system to provide simultaneous sampling over 32 channels, 24-bit resolution, 10 kHz bandwidth, 50 kHz sampling rate, and battery-powered operation while reducing cost by two orders of magnitude. System performance was evaluated comparatively against a reference device. Initial validation involved benchtop recordings in saline solution and standard non-invasive biopotential measurements (ECG and EMG). Further in-vivo validation was performed by recording evoked electrophysiological responses and FN-EIT traces from the sciatic nerve of a rat during tibial branch stimulation.

**Main results:** EMG recordings showed comparable RMS peak amplitudes (814±153 µV vs. 897±113µV, p=0.07), while ECG-derived heart rates closely matched between systems (64.8±4.0 bpm vs. 65.1±3.1bpm, p=0.54). During in-vivo recordings, compound action potentials exhibited comparable amplitudes and morphology (129±26 mV vs 128±25 mV, P=0.15). FN-EIT recordings showed strongly correlated baseline voltages (R>0.93, P=0.11), sub-microvolt noise levels (0.83±0.36µV vs. 0.42±0.25µV, p<0.05), and comparable impedance variations (0.006±0.002% vs 0.005±0.003, P>0.05). FN-EIT images of functional activity recorded with the novel device closely matched reference ones, exhibiting a 98.5% overlap in activated area.

**Significance:** The proposed open-source device has the potential to broaden research access to customizable, high-specification data acquisition hardware and facilitate wider adoption of specialized neural recording techniques such as FN-EIT.

## 1. Introduction

Biopotentials are electrical signals generated by the physiological activity of excitable cells, primarily neurons and muscle fibers. The acquisition and analysis of biopotentials play a fundamental role in medicine, enabling monitoring of a wide range of physiological and diagnosis of pathological processes. Among non-invasive measurement techniques, electrocardiography (ECG), electroencephalography (EEG), and electromyography (EMG) are the most widely used [1]. These modalities record electrical activity from the heart, brain, and skeletal muscles, respectively, typically using surface electrodes placed on the skin. In addition to surface measurements, biopotentials can also be recorded invasively at the neural tissue level, providing higher spatial and temporal resolution and enabling direct observation of neural activity. Examples include electrocorticography (ECog)[2] and local field potentials (LFPs)[3] in brain, and compound action potentials (CAPs) in peripheral nerves [4]. These signals are typically acquired using implanted electrodes placed in close proximity to neural structures, allowing the measurement of electrical activity from populations of neurons or even individual cells, such as in the case of single-unite recordings [5].

Biopotential measurement hardware provides the interface between physiological electrical activity and digital systems used for visualization, storage, and analysis. Regardless of the target signal, the functional architecture and main building blocks typically follow a similar organization [6]. A standard biopotential acquisition chain consists of electrodes, protection circuitry, a high-input-impedance instrumentation amplifier, filtering stages, gain control, and an analog-to-digital converter (ADC). A digital logic block, such as a microcontroller, DSP or FPGA is used for device control and data transmission. This architecture is largely shared across different applications, with variations primarily reflecting the characteristics of the signal being measured, particularly in parameters such as gain, frequency bandwidth, and input dynamic range. Design implementations generally fall into two categories: application-specific integrated circuits (ASICs) and systems based on off-the-shelf components. ASIC solutions integrate multiple functional stages within a single chip, offering advantages in power consumption, compactness, and channel density, which are particularly beneficial for wearable devices and high-channel-count systems. In contrast, off-the-shelf solutions based on discrete commercial components provide greater flexibility and enable faster prototyping, while avoiding the substantial cost and time investment required to design and manufacture custom ASICs. Another relevant distinction is between clinical-grade and research-grade systems. Clinical devices are typically designed to meet the minimum ADC vertical resolution and bandwidth recommended by guidelines for a given bioelectric signal. In contrast, research-grade systems generally provide higher vertical resolution, wider bandwidth, lower noise, and the highest possible SNR in order to maximize the detection of very small signal components with potential scientific relevance. A typical example of this distinction is the use of 16-bit ADCs vs. 24-bit ADCs.

In addition to established commercial options for recording biopotentials, open-source hardware also exist, typically developed for student education, research applications, or deployment in low-resource settings; among those, many recent projects [7–10], tend to rely on off-the-shelf components integrating an Analog Front-End (AFE) and an ADC within the same IC, such as Texas Instrument’s ADS1298 and ADS1299.

Recently, the novel technique known as Fast Neural Electrical Impedance Tomography (FN-EIT) has also emerged as a promising method for monitoring neural activity in the brain and peripheral nerves (Fig. 1). Electrical Impedance Tomography (EIT) has been originally investigated for thoracic imaging [11], before expanding to applications across a variety of biological tissues [12]. Unlike conventional electrophysiological recordings that measure extracellular biopotentials, Fast Neural EIT measures transient changes in the bulk electrical conductivity of neural tissue, generated by the opening and closing of ion channels during action potentials [13]. A set of electrical impedance measurements collected from multiple electrodes over time is then subject to image reconstruction for localising the source of neural activity [14]. Fast Neural EIT can achieve measurements of neural activity with high temporal resolution, in the order of milliseconds; however, a highly sensitive data acquisition system is required which is capable of detecting extremely small impedance changes. In the majority of cases, Fast Neural EIT is performed using the ScouseTom recording system [15], a modular EIT platform integrating commercial devices for current injection and signal acquisition with custom circuitry responsible for multiplexing and digital control. While the system incorporates several commercial modules, a fully open-source implementation would be preferable. Such an approach would facilitate broader dissemination within the research community and enable adoption by laboratories with varying levels of funding and resources. Previous work already demonstrated the feasibility of replacing the benchtop currents sources employed for neural tissue stimulation and impedance measurement with custom circuitry [16]. However, a research-grade EEG system is currently used for acquiring data with a digitization resolution of 24-bit resolution and a significant level of oversampling, both necessary to achieve a Signal-to-Noise Ratio (SNR) sufficient for the reliable extraction of impedance changes associated with neural activity. To date, no custom open-source module has been developed that enables data acquisition specifically for FN-EIT.

**Figure 1.**
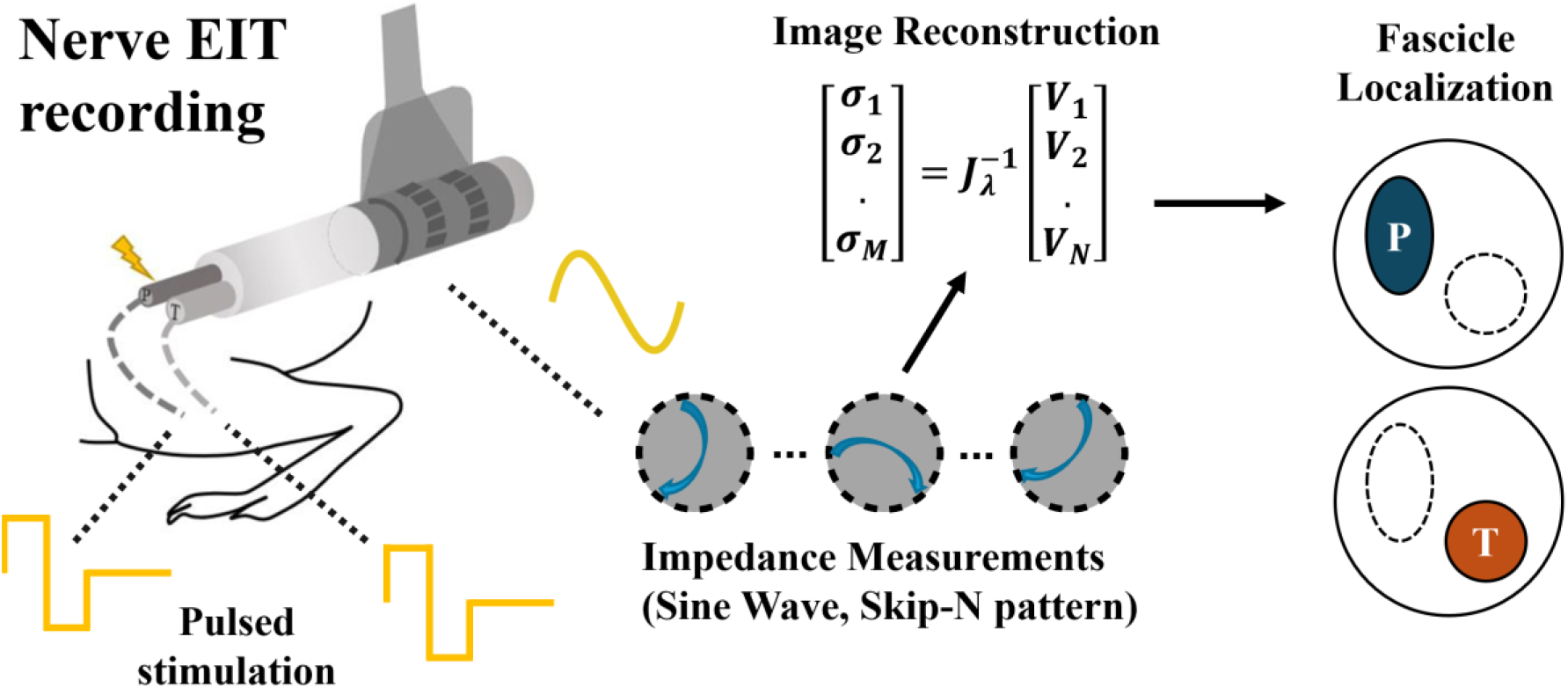
Visual summary of the established fast neural EIT nerve experiment protocol. Pulsed current is delivered to individual nerve branches, such as Tibial (T) and Peroneal (P) branches in the sciatic nerve. Multiple impedance measurements are performed with sinusoidal current on selected electrode patterns within a circular nerve cuff electrode array. Impedance data is fed to a reconstruction algorithm to localise the activated fascicle within the cross-section of the nerve.

In summary, we identified the need for a low-cost biopotential recording system with high-end features comparable to those of research-grade systems, while remaining open-source and also potentially suitable for FN-EIT applications. With this project, we aimed to:

i. Deliver an open-source, compact, high-end biopotential amplifier with multiple use cases and relatively low-cost compared to commercial options with similar specifications.
ii. Demonstrate successful use in common biopotential recording applications such as recording ECG, EMG and EEG signals.
iii. Demonstrate successful use in more specialised applications such as invasive electrophysiological recordings and fast neural EIT imaging.

## 2. Study Design and Outline

Our project started with component selection and electronic board design. We aimed to design a recording device fitting the following specifications:

- Only off-the-shelf components available from main IC suppliers, no custom ASICs or uncommon ICs.
- Analog input range ≥400mV, to support EIT applications
- ADC resolution 24 bit
- Minimum of 32 analog input channels, simultaneous sampling, individual channel sampling speed ≥50kHz
- Bandwidth ≥10kHz
- Presence of auxiliary channels for trigger detection and synchronization
- Noise levels comparable to similar high-end amplifiers (max 50% or 5µV higher, whichever is lowest)
- Analog front-end should be isolated and battery-powered for electrical safety and reduction of mains supply noise.
- Device should be complete with a digital control system (e.g. microcontroller or FPGA) and USB interface to PC.
- Price, including manufacturing and component cost, in the order of a few hundred GPB, ideally <500USD.

During the early phases of the project we went through multiple iterations, progressively improving functionality and reducing size of the device until reaching the final version presented in this manuscript and satisfying the specifications above. The device then underwent multiple testing phases.

We first developed a calibration process by feeding test signals to the device in order to estimate DC offsets and gain mismatch between different channels. The resulting calibration matrix was applied to each subsequent recording to improve accuracy. Then, we recorded data from a nerve cuff suspended in saline solution to assess basic noise levels in the frequency band of biopotential and electrophysiological signals, and in the frequency band typically used for neural EIT, while injecting sinusoidal waveforms. Following the establishment of a calibration protocol and acceptable noise levels, we performed a series of exemplary recordings to assess the capability of our system to collect common non-invasive biopotential data such as ECG, EMG and EEG signals on live subjects. The final and most complex test we conducted involved collecting in-vivo electrophysiological data and fast neural EIT recordings from the peripheral nerve of a live animal using a multi-electrode nerve cuff, effectively replicating previous experiments from our group but with the novel device.

All “wet” recordings performed in either saline solution, live subjects and animals were either preceded or followed by recordings performed in identical conditions with a reference commercial amplifier.

## 3. Methods

### 3.1. Device Architecture

A block diagram which includes the main stages of electronic architecture is shown in Figure 2. The core component for any data acquisition device is the ADC. For this project, we chose the AD7771 (Analog Devices). This eight-channels ADC satisfies our requirements of having 24bit resolution, high sampling rates and simultaneous sampling. It also features digitally selectable input gain, which adds flexibility toward different applications, and Electroencephalography is specifically mentioned in the datasheet as an application. Four identical ADCs were used, all connected to the same master clock oscillator operating at 8.192 MHz. Communication with the ADCs for the purpose of setting AD conversion parameters and data transfer was handled through SPI protocol. SPI pins were shared between ADCs, except for the pin necessary for output data transfer toward the digital control logic. Synchronization of operations across the four ADCs was addressed as suggested by the datasheet, i.e. a synchronization pulse between ICs. A +2.5V voltage reference (ADR441, Analog Devices) was connected to all the ADCs.

**Figure 2.**
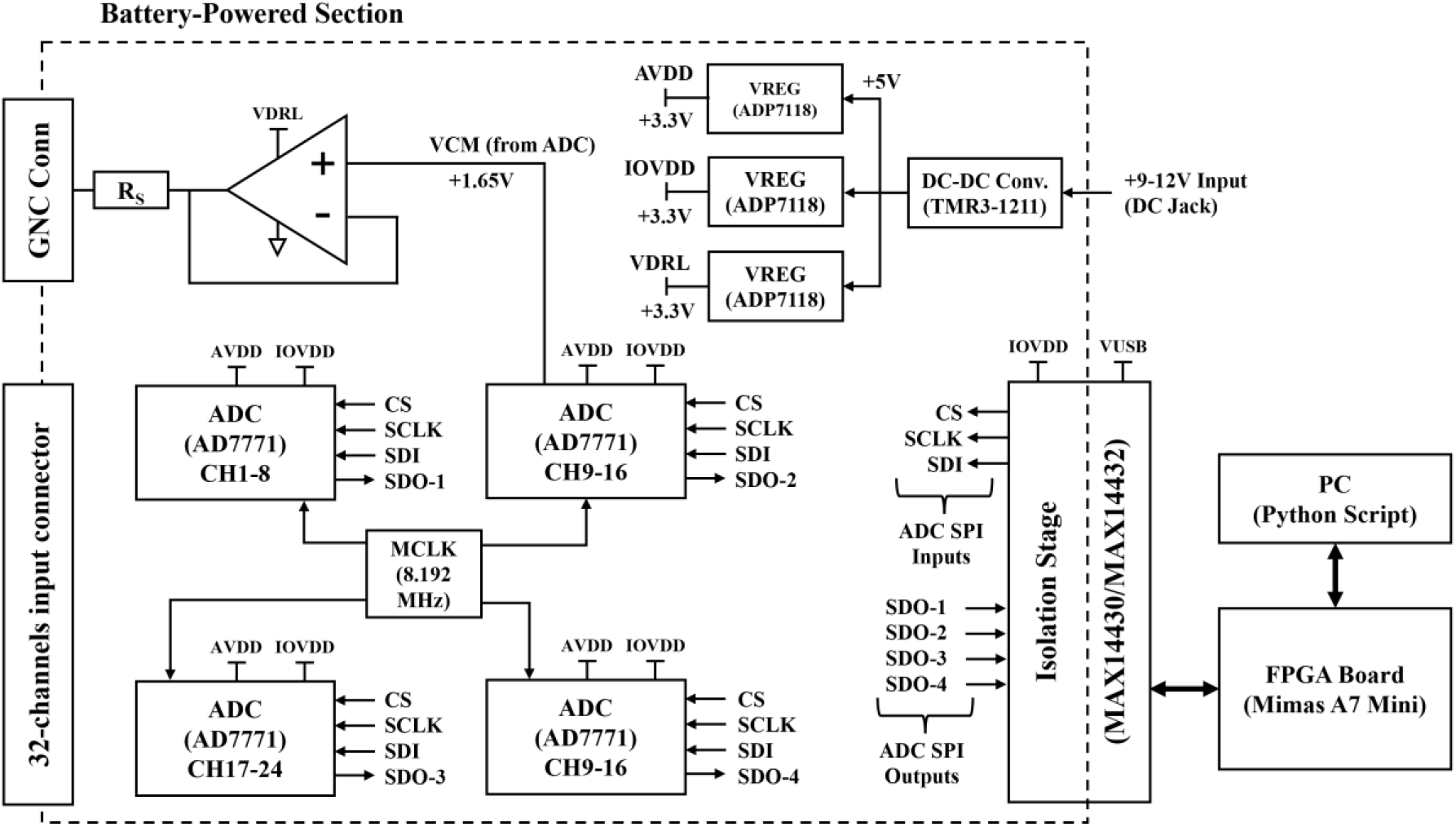
Block diagram of the novel open-source EEG system. Four 8-channel AD7771 collect and digitise input data through shared SPI lines. Electrical insulation is ensured between the front-end and the FPGA board through MAX14430/32. An operational amplifier-based circuit applies mid-point bias to the tissue. Separate power regulators are used for powering analog, digital and bias circuitry.

Our chosen ADC integrates a programmable gain amplifier and ESD protection on each channel, reducing the amount of required external circuitry. However, additional circuitry is needed to ensure that input signals fall within the ADC input range, i.e. inputs should be biased around the ADC common-mode voltage. Typically, one of two main schemes for bias compensation is used:

1. The Driven Right Leg (DRL) scheme [17]
2. Differential output ADC drivers [18]

The main drawback of scheme #1 is that two specific channels must be selected to extract common-mode voltage of the subject or biological tissue. This can be problematic in systems such as nerve electrode cuffs, where some electrodes may not be in contact with the tissue or may suffer from faulty interconnects.

Although most modern systems allow users to specify which input channels participate in the DRL loop, this still requires a manual channel-selection procedure. Scheme #2, however, requires a substantial number of additional external components for each input channel. In designing our device, we sought a bias compensation mechanism that minimizes component count and operates independently of channel selection. For this reason, we adopted the more recent “DRL-less” approach [19]. In our final configuration, all ADC positive input pins are accessible through the input connector, while the negative input pins are all connected to the ADC common-mode pin V_cm_. V_cm_ is also fed to an operational amplifier in buffer configuration (IC) whose output is connected to the subject or biological tissue. A 10kΩ series resistor is applied to the buffer to improve electrical safety. Thus, all data acquisition is performed in respect to the ADC V_cm_ value, while the body is biased to be as close as possible to V_cm_. Following data acquisition, a reference channel picked by the used is digitally subtracted from all the other input channels. Notch filtering of mains supply is also performed in software.

Power supply and regulation in the device is organized in multiple stages. The ADCs, master clock oscillator, power-on LED and bias circuitry are battery-powered, whereas digital control circuitry is powered directly through its USB connection. A DC barrel jack acts as input connector for the battery; any 9-12V input is acceptable, such as 9-12V power banks, common 9V block batteries, or larger, higher capacity batteries. The power input is converted down to 5V by a DC-DC converter (TMR3-1211, Traco Power) and then converted to +3.3V through three separate voltage regulators (ADP7118, analog Devices, Fixed 3.3V ouput version). The three identical 3.3V regulators supply the bias circuitry and the ADC analog and digital power input pins, respectively.

Digital communication between the ADCs and the digital control circuitry is routed through digital galvanic isolator. High-speed isolators were chosen due to the fast transmission speeds involved (MAX14430/MAX14432, Analog Devices). A total of 3 isolators were used.

Digital control and data transfer from the ADCs to PC is handled in the final device through a commercial FPGA board (Mimas A7 Mini, Numato Labs) hosting an Artix-7 FPGA chip. This specific board was chosen for its small size compared to other FPGA development boards, and for the presence of an embedded high-speed FIFO-to-USB2.0 IC (FT2232H, FTDI), which allows both firmware upload and data transfer on separate channels. USB2.0, also known as high-speed USB, was necessary to handle the data transfer rate from 32 channels operating at 24bit and 50kHz sampling rate. Design of a custom FPGA board was considered out of scope for this project. FPGA Firmware was written in Verilog and developed on the VIVADO IDE (2024 Edition, AMD). It features an initialization sequence to set up desired ADC conversion gain and sampling speed and then stabilizes into a conversion and data transfer loop based on start and stop commands from the connected PC. Data bytes coming from the ADCs and auxiliary channels are collected into a data frame and pass through a custom FIFO buffer before streaming to PC through the USB chip. On PC, a Python script collects incoming data and stores it into raw binary format. A MATLAB script is used for readout and conversion from the binary file; further signal analysis in this study was also performed in MATLAB.

In terms of physical layout, the device was implemented using three vertically stacked Printed Circuit Boards (PCBs) to achieve a compact form factor and maintain balanced dimensions. This approach avoided excessive elongation along any single axis, which could otherwise result in an impractical design. The top PCB hosts the first stage of voltage regulation, the output bias circuit with its own +3.3V voltage regulator and the 32-channel input connector. The middle PCB hosts the four ADCs, the local clock source, and the +3.3V voltage regulators for digital and analog ADC power supply. The bottom PCB is dedicated to digital control and includes a socket for the FPGA module, the digital isolators, and pushbuttons connected to the FPGA input pins. PCBs were designed with 4, 6 and 2 copper layers respectively from top to bottom. See Data Availability section for access to schematics and firmware.

### 3.2. Basic Testing and Calibration

Our first step for evaluating device functionality was to deliver a set of constant voltages to all the device input channels simultaneously. The set of voltages was generated using a KeySight 33600A Waveform Generator (KeySight, Santa Rosa, USA) and consisted of 25 voltages equally spaced in 50mV steps in the range of ±600mV. Accurate estimates of the true voltage values were collected using a benchtop HP34401A voltage meter (Agilent Technologies, Santa Clara, USA).

In addition to confirming basic functionality, these measurements were also used to perform calibration of individual channels through linear regression between values measured by the multimeter and the device. The set of calibration coefficients determined by this method were stored externally to the instrument and applied to all data collected in the rest of the study.

### 3.3. Noise Levels in Saline Solution

We first evaluated the novel device in wet conditions by placing a Multi-Electrode Array (MEA) nerve cuff, designed for recordings in peripheral nerves of large animals [20] within a beaker filled with saline solution (Fig.3-a). The solution was prepared as a mixture of deionized water and Sodium Chloride (NaCl), with concentration adjusted to reach electrical conductivity ≈0.3 S/m, representative of typical bulk conductivity of neural tissue [21–23].

**Figure 3.**
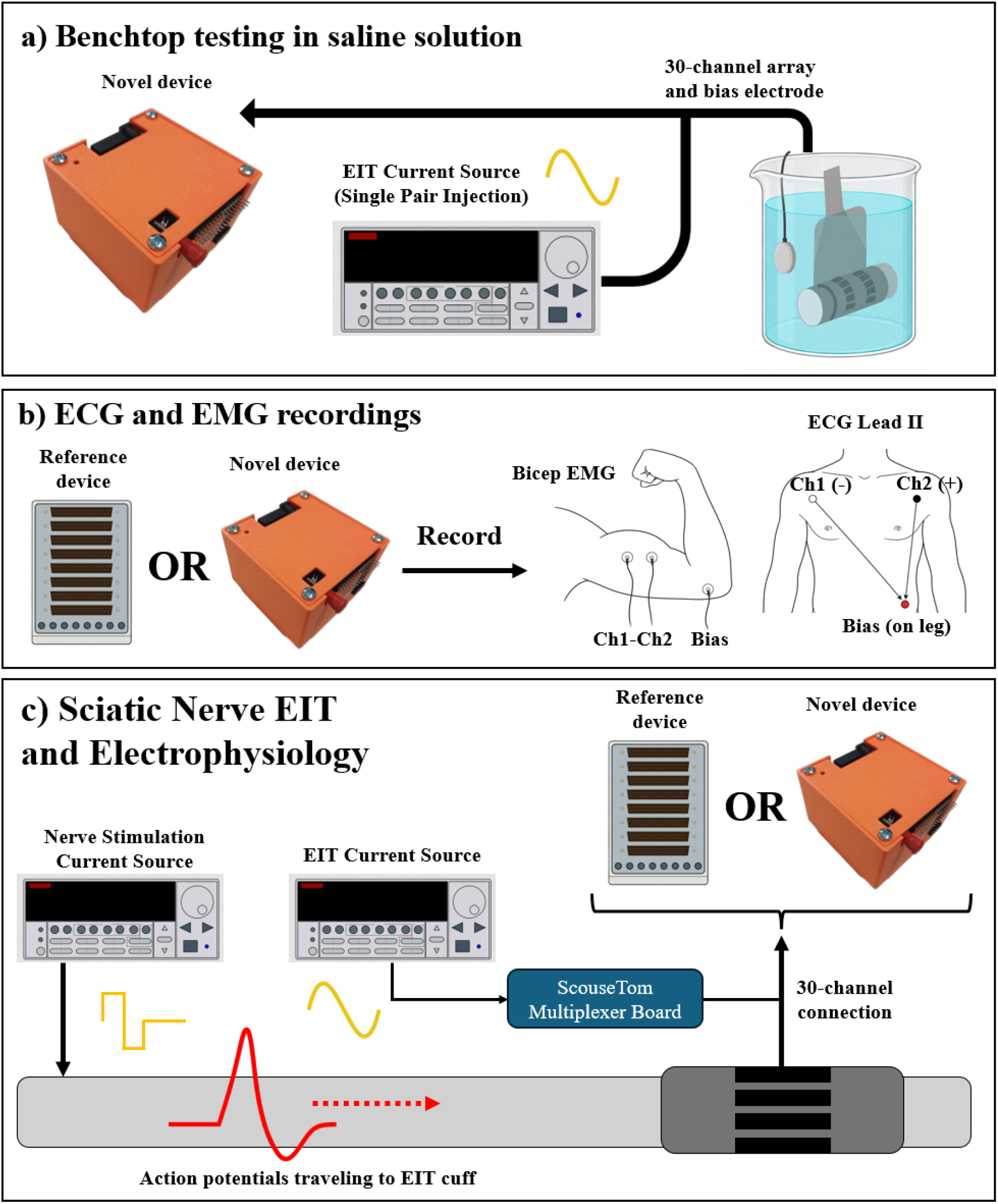
Summary of experimental work performed in this study. a) A nerve cuff array is placed in diluted saline solution to assess noise levels of the device. b) ECG and EMG recording are performed with both the novel and a reference device for comparison. c) Nerve electrophysiological and FN-EIT recordings are performed on rat sciatic nerve using a full ScouseTom EIT system, with either the reference or novel device acting as a biopotential amplifier.

In these conditions, we first performed recordings with no input, to assess baseline noise levels, and then performed recordings while injecting AC sinusoidal current at 100 µA and 6 kHz using a previously designed AC current source [16]. Resulting data was demodulated within 1kHz bandwidth of the carrier frequency using the Hilbert transform to assess noise levels in fast neural EIT application. Control recordings were performed in comparison with the same commercial EEG amplifier used throughout the rest of the project as a gold standard (Actichamp EEG recorder, Brain products GmbH, Germany), named from now on as the reference device.

Power consumption was also assessed at this stage by measuring voltage drop on a small series resistor embedded in the power supply cable.

### 3.4. EMG and ECG Data Acquisition

EMG recordings were performed in a standard configuration for detecting bicep contraction, by placing two electrodes centered on the muscle belly at 2cm distance from each other and the ground electrode opposite in the elbow area (Fig.3-b). A simple experiment was conducted in which the subject’s bicep was alternately contracted and relaxed for 5 seconds each over a few minutes. ECG recordings were performed in a standard configuration for recording the lead II signal (Fig.3-b), by placing recording electrodes on the right arm and left leg, with the ground electrode on the right leg. ECG signal was collected at rest from a test subject over a few minutes. Each type of recording was performed with both the proposed device and the reference device.

Collected data was subject to the same signal processing steps regardless of data acquisition system. ECG data was subject to bandpass filtering in the 0.05-150 Hz bandwidth. Then, HR was detected by finding local peaks in the ECG signal. EMG data was subject to bandpass filtering in the 10-500 Hz bandwidth and RMS envelope detection with a low-pass filter cut-off frequency of 2 Hz. A 50Hz digital notch filter was applied to all recordings.

### 3.5. Electrophysiology and Neural EIT Data Acquisition

We performed a representative peripheral nerve experiment to prove the feasibility of using our device to collect in-vivo electrophysiological and FN-EIT data. The overall experimental setup and protocol also followed from previous work on nerve EIT, as summarised in Fig. 1, with the difference that recordings were performed with both the original setup (ScouseTom) and the novel device (Fig.3-c).

The experiment was performed on the sciatic nerve from an adult male Sprague-Dawley rat weighing ≈450g, with the same surgical and anesthesiologic procedures as in previous work [21– 23]. All animal experiments undertaken in this study were approved by the UK Home Office and in accordance with its regulations.

Briefly, a previously described nerve EIT MEA cuff [24] was placed around the main trunk of the sciatic nerve, and a smaller stimulation cuff electrodes (CorTec Gmbh, Freiburg, Germany) was placed around the tibial fascicle at ~1.5 cm distally from the EIT cuff. The EIT cuff consisted of 14 electrodes arranged in a ring-like arrangement and was manufactured with stainless steel and silicone polymer and coated with a conductive polymer to improve electrochemical properties [20]. Fascicle stimulation was performed by delivering biphasic 50 µs pulses at 2mA amplitude and 200 ms repetition time though each individual stimulation cuff. EIT current was injected at 150µA and 6 KHz as in previous work [24]. Fourteen injection electrode pairs were used in a skip-4 configuration; 30s were spent recording data from each injection pair, leading to a total of 7min recording time for each fascicle. Raw signals were converted to EIT traces by performing band-pass filtering at 1KHz bandwidth around the 6 KHz EIT carrier and then performing Hilbert transform demodulation (modulus only) for the purpose of envelope extraction. Coherent averaging over repeated stimulation pulses was performed to reduce background noise. Electrical stimulation of the individual fascicular branches was also performed before and after EIT recordings to capture standard electrophysiological markers at the levels of the EIT cuffs; specifically, Compound Action Potentials (CAPs) were recorded from the EIT cuff, and their presence was considered an indication of nerve tissue health throughout the experiments.

All EIT and electrophysiology measurements were performed with a ScouseTom system [15], which features a commercial benchtop current source (Keithley 6221, Tektronix,USA), a commercial high-end EEG amplifier (Actichamp, Brain products GmbH, Germany), and custom electronics for digital synchronization and current source electrode switching. For the purpose of our experiment, each recording was performed twice, once with the full ScouseTom setup and once with the EEG amplifier replaced by our novel device. Due to the different cut-off frequencies of the input anti-aliasing filters in the reference and custom amplifiers (7.5 kHz and 10.5 kHz, respectively), a frequency response–based correction factor was applied to the reference recordings to enable amplitude comparison with the custom device. The frequency responses of both devices were measured in the 1-10kHz range in 1 kHz increments, and the corresponding relative attenuation factors at 2 kHz and 6 kHz were applied to the CAP and EIT data, respectively.

After being subject to the same outlier rejection criteria previously adopted [21], EIT δV traces were used as input for an image reconstruction algorithm [24] to identify areas of peak fascicular activation over the cross section of the nerve. Briefly, a 1.8M-elements forward model of the nerve was built in the EIDORS Matlab plug-in [25] and used to generate a Jacobian sensitivity matrix for our EIT protocol [24]. A coarse voxel-based version of the Jacobian matrix and original mesh were generated with voxel size 40 μm; mathematical inversion to estimate conductivity distribution was performed using 0th-order Tikhonov regularization with noise-based voxel correction [26]. The resulting 3D distribution of relative conductivity variation δσ was sliced across the area beneath the electrode positions to generate images of fascicular activation over the cross-section of the nerve. Comparison between standard ScouseTom configuration and the novel device was performed on both signals and reconstructed images. For CAP signals and EIT δV traces, baseline noise amplitude, peak signal amplitude, peak timing and SNR were compared. For reconstructed images, we calculated the percentage of overlap between areas of peak fascicular activation reconstructed from both hardware configurations and the cartesian distance between the Centers-of-Mass (CoMs) of the reconstructed images.

## 4. Results

### 4.1. Board Manufacturing

The final device iteration had a compact size of ≈85×85×60mm, power bank excluded and is shown in Fig. 4 with and without a custom 3D-printed protective case. At the time of manufacturing (June 2025), the total price of PCB production component purchase and assembly was ≈336 USD for a single device (FPGA board excluded), two orders of magnitude lower than commercial high-end EEG recorders. More specifically, total cost for manufacturing the 3 PCBs was 26 USD, component cost ≈210 USD and assembly was ≈100 USD. Overall, cost was mostly concentrated in the middle board (253 USD for components and assembly), which has six copper layers, hosts the most expensive components such as the four high-end ADCs, and required assembly for the largest amount of SMD components.

**Figure 4.**
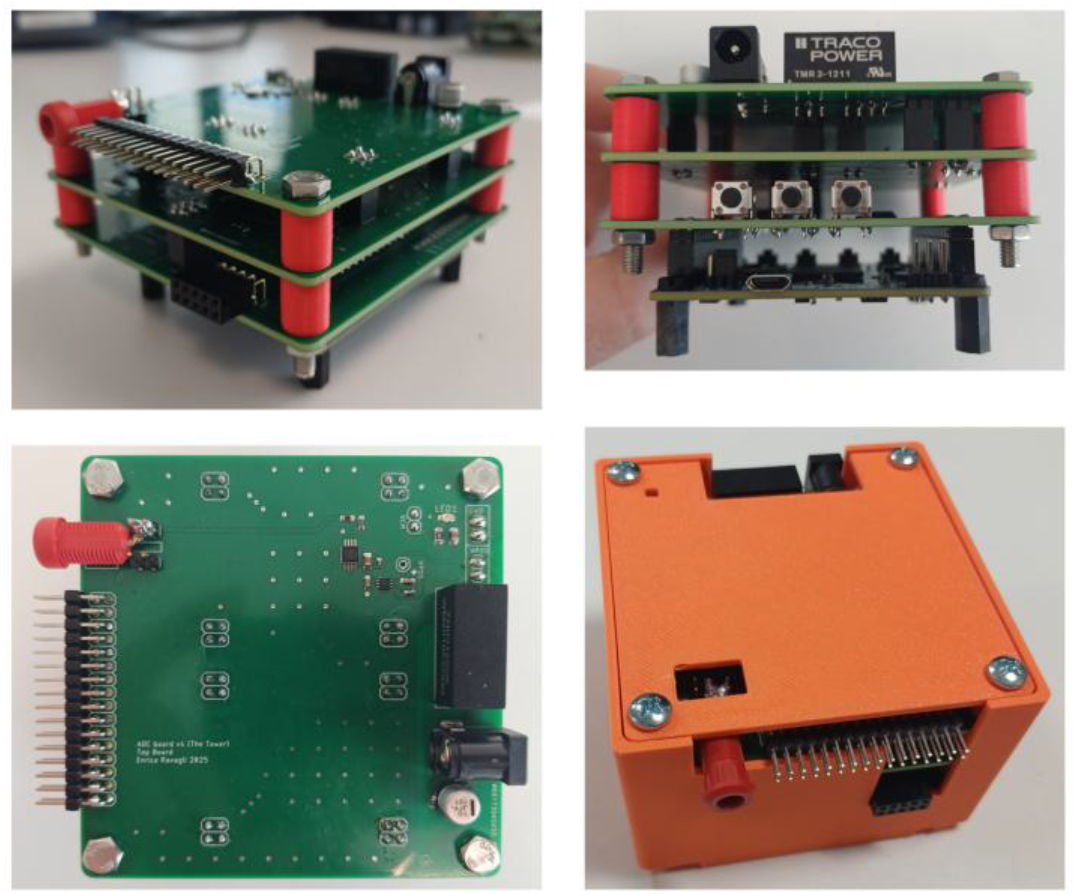
The novel biopotential amplifier. The device is composed of three custom stacked PCBs and an FPGA board connected to the bottom layer.

### 4.2. Basic Testing and Calibration

Linear regression coefficient computed for each of the 32 input channels of the novel device were 1.0007 ±0.0002 mV and 1.5368 ± 0.0118 mV for gain and offset, respectively.

Pre-calibration RMSE, evaluated on all data points from all channels in a range of ±600mV, was 1.5697mV; post-calibration, in the same conditions, it was 0.2251 mV. Correlation between measurements performed by the novel device and the multimeter was R>0.99.

A correlation plot between the reference and measured values post-calibration is shown in Fig. 5-A. A Bland-Altman plot of the difference between values measured by the ADC and the reference instrument in shown in Figs. 5-B and 5-C for pre- and post-calibration conditions, respectively.

**Figure 5.**
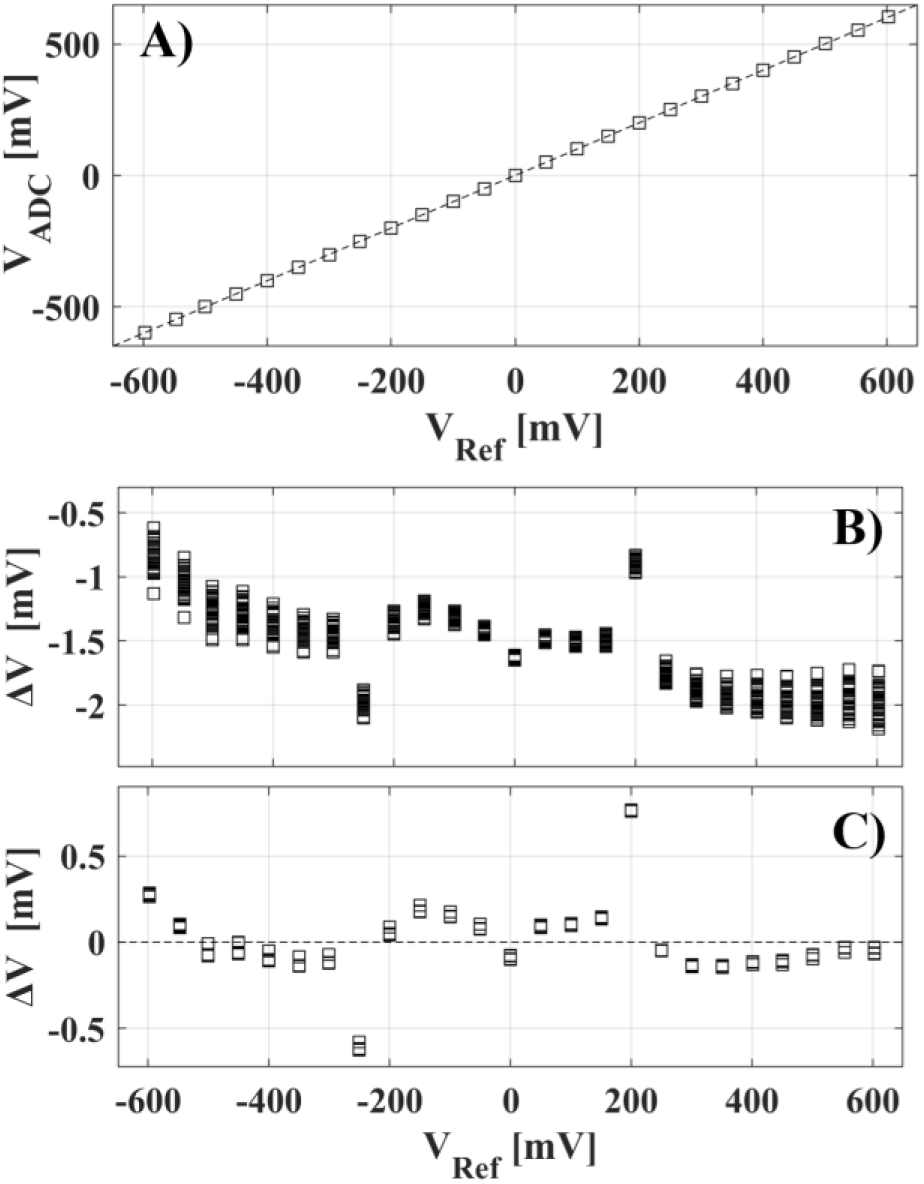
A) Correlation plot between voltages acquired by the reference and novel device, post-calibration. B-C) Bland-Altman error plots for values measured by the novel ADC device in pre- and post-calibration conditions, respectively.

### 4.3. Noise Levels and Power Consumption

In saline solution, average baseline noise levels in the 0.1Hz-10kHz bandwidth over N=29 electrodes were 12.4±0.3µV and 10.5±0.8µV (p=4.5174e-10) for the custom and commercial devices, respectively. Noise levels within a 1kHz bandwidth of the demodulated EIT carrier frequency were 3.5±1.3µV and 3.8±0.7µV (P=0.1959) for the custom and commercial devices, respectively. Power consumption during recordings was 158mA, enabling ≈57 hours of operation from a 9000mAh 9-12V power bank.

### 4.4. EMG and ECG Data Acquisition

Our novel device was successfully able to collect EMG and ECG data visually and numerically comparable to data acquired with the reference instrument. Examples of EMG signals acquired with reference and novel devices are shown in Figs. 6-A and 6-B with and without envelope extraction. RMS-EMG peak values averaged over n=18 repeated contractions were 814±153µV and 897±113 µV (p= 0.07) for the reference and novel device, respectively.

**Figure 6.**
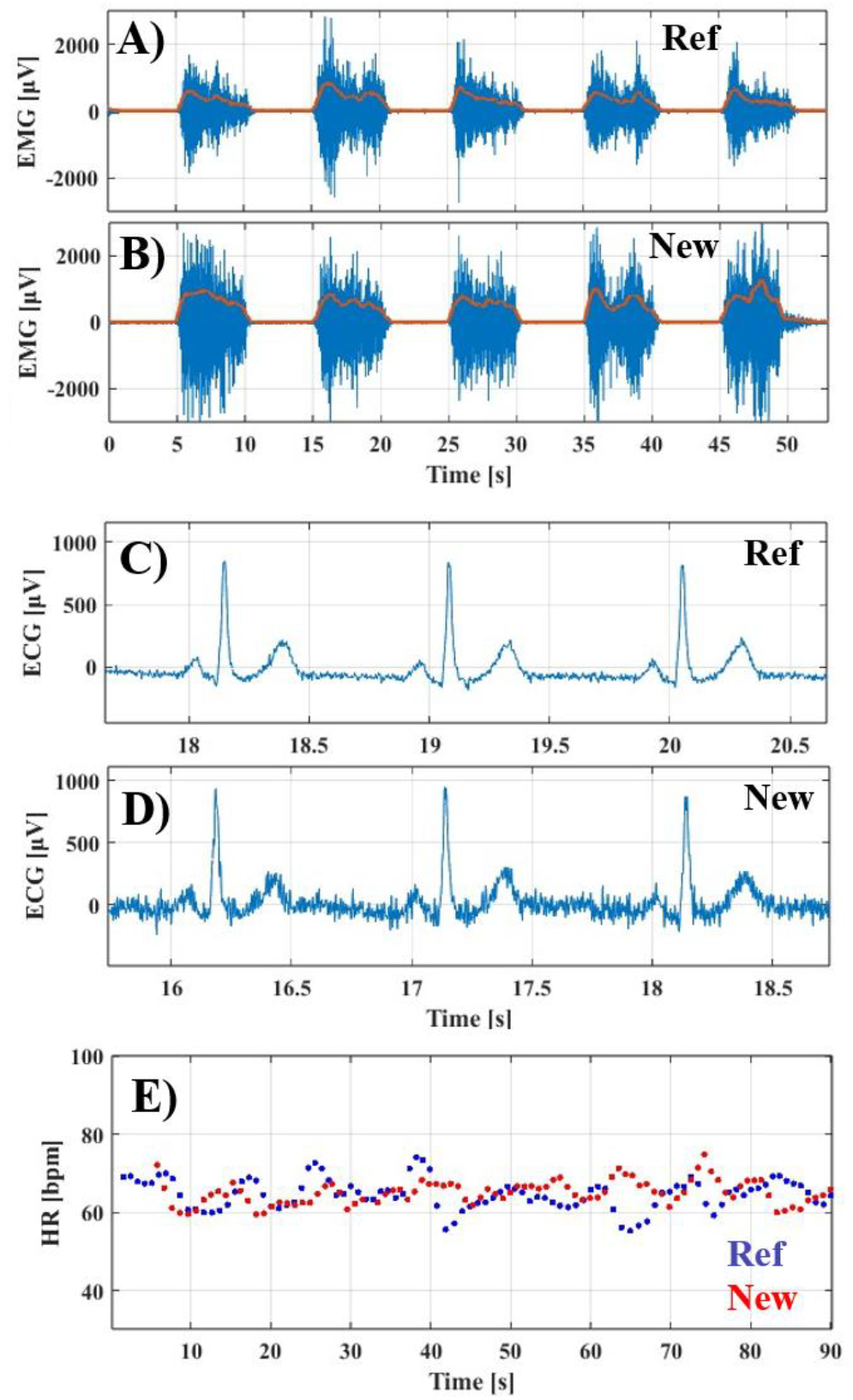
A-B) EMG recordings performed with the reference and novel devices. Blue and orange lines represent EMG signals before and after RMS conversion, respectively. C-D) ECG recordings performed with the reference and novel devices. E) Heart rate extracted from ECG recordings.

Exemplary ECG Lead-II traces acquired with reference and novel devices are reported in Figs. 5-C and 5-D and show consistent detection of major ECG signal components between devices, with marginally higher noise levels in the custom device. Heart rate data extracted from R-R peaks in ECG data is shown in Fig. 6-E. Resting HR values averaged over n=91 heartbeat events were 64.8±4.0 bpm and 65.1±3.1 bpm (p=0.54) for the reference and novel device, respectively.

### 4.5. Electrophysiology and Neural EIT Data Acquisition

Data collected in-vivo from N=1 animal with neural activity evoked by stimulation of the tibial fascicle returned overall comparable results between the novel and reference devices after frequency response-based correction (Fig. 7). CAPs collected by the novel device had comparable peak amplitudes to the reference (129±26 mV vs 128±25 mV, P=0.15) and displayed visually similar waveform shapes (Fig.7a-b). Baseline EIT voltages (BVs) collected by custom and reference devices were highly correlated (R>0.93, P=0.11). EIT data collected within the experiment was averaged over n=150 repeated events, corresponding to 15s of fascicular stimulation at 10Hz, for each injection pair.

**Figure 7.**
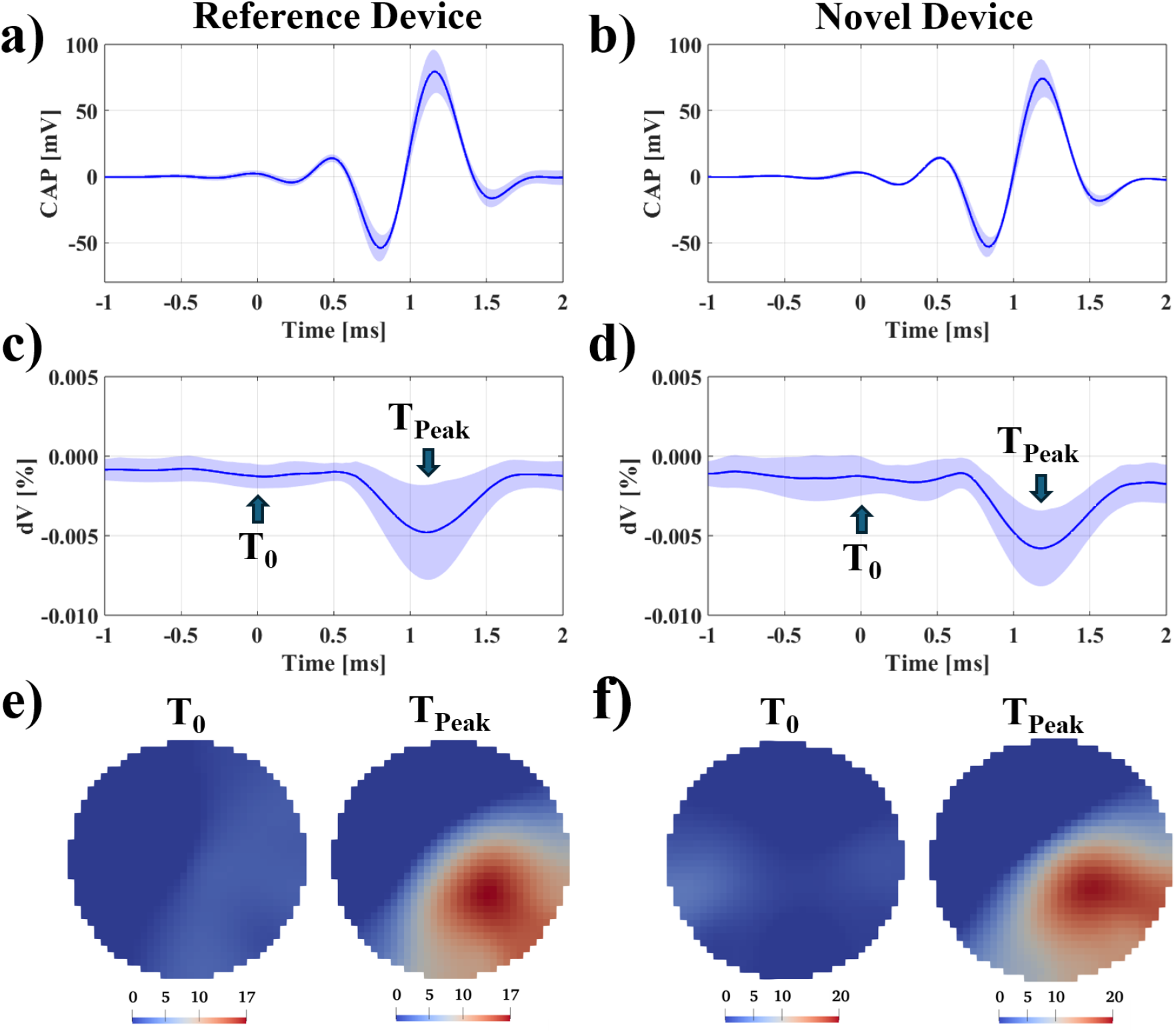
a-b) Compound action potentials recorded from rat sciatic nerve cuff electrodes with reference and novel device in response to stimulation of the tibial fascicle at time t=0. c-d) EIT δV traces with BV>50mV collected with the same stimulation parameters as a-b). Mean is represented as blue continuous line and shaded areas representing standard deviation. All data in a-b-c-d) is averaged over 150 events. e-f) Image reconstruction of fascicular functional activation over the cross-section of the nerve at baseline and peak activation time, performed from data recorded from both devices.

Post-averaging, noise levels in the nerve EIT bandwidth were 0.83±0.36µV vs 0.42±0.25µV (P<0.05) for the novel and reference devices, respectively. Peak amplitude of evoked δV neural impedance changes detected by both devices (Fig.7c-d) was comparable in amplitude but showed statistically significant difference (3.2±1.7 µV vs 2.4±1.6 µV, P<0.05). Combined, noise and peak δV results led to a peak SNR performance of 4.8±4.1 vs 7.6±7.2 for the novel and reference devices, within this particular nerve. When expressed as percentage change relative to baseline voltage, peak δV% values were comparable between the novel and reference devices (0.006 ± 0.002% vs. 0.005 ± 0.003%, P > 0.05). For this analysis, δV traces associated with BVs<50mV were excluded to prevent disproportionate amplification of noise arising from normalization by small baseline values.

Image reconstruction was performed on δV values at peak time from recordings obtained with both devices (Fig.7e-f). The resulting images revealed clear areas of functional activation across the nerve cross-section, which were strongly co-localized between recordings. Overlap between activation areas in images reconstructed from both devices, defined as pixels with at least 50% of peak intensity, was 98.5%. Distance between CoMs calculated on the same pixel groups was 9.6µm, lower than the size of a single pixel in our mesh, or 0.7% of the nerve diameter.

## 5. Discussion

Within this project, we developed a complete open-source solution for high-speed, high-resolution acquisition of electrophysiological data in a compact format. The low cost of the proposed device enables implementation of a high-performance bioelectrical recording system at a fraction of the cost of conventional high-end platforms, improving accessibility for research laboratories. Although the reported cost excludes the selected FPGA board, which adds approximately USD 150 to the total system cost, the FPGA is a reusable general-purpose platform that can support multiple laboratory projects rather than being a cost solely allocated to this device.

### 5.1. Comments on Experimental Results

We evaluated our device across a diverse set of applications to thoroughly assess its general performance. EEG was not recorded for this study, but the successful measurement of ECG, EMG, and even neural activity at the surface of the nerve suggests that EEG acquisition would also be completely feasible. ECG recordings exhibited a slightly elevated noise baseline compared to the reference device, which could likely be mitigated through improved or dedicated filtering. Nonetheless, the waveform morphology remained clearly discernible, allowing comparable detection of ECG features and heart rate.

Our device successfully recorded invasive signals from a nerve cuff electrode placed on the rat sciatic nerve. The amplitudes and shapes of compound action potentials (CAPs) and baseline voltages (BVs) were consistent with those obtained using the reference device. Although δV peak values differed slightly in absolute terms, they were identical when normalized to baseline voltages, suggesting that discrepancies in absolute values likely reflect physiological changes in the nerve over the course of the experiment, which affected the amplitude of recorded impedance changes. Noise level in EIT bandwidth, while higher for the novel device, remained below the typical ≈1 µV threshold generally considered acceptable for successful FN-EIT recordings, indicating that the new device is suitable for use with this technique. This conclusion is further supported by the images reconstructed from δV traces, which appear nearly identical across devices, with closely matching CoMs and overlapping activation regions.

Compared to our previously reported work, where average recorded δV peaks where in the range of 10–38 µV [21,22], the response in this animal was much lower. This outcome demonstrates that our system is capable of detecting impedance changes and reconstructing corresponding images even under low-signal conditions.

### 5.2. Limitations

While the proposed system serves its purpose in filling a gap within the set of available open-source research instruments, the present implementation is subject to certain limitations.

The current system supports simultaneous sampling of 32 channels up to 50 kHz. In contrast, comparable commercial platforms provide sampling rates of up to 100 kHz across 32 channels, or 50 kHz across 64 channels. Future iterations will focus on increasing both the maximum sampling rate and the number of available input channels.

The current version provides eight digital input pins for auxiliary signals such as triggers; however, it does not include analog auxiliary inputs for recording signals such as temperature or pressure. Future iterations will incorporate a set of analog auxiliary inputs that are electrically isolated from the primary input channels.

Increasing the sampling rate and/or the number of channels will require higher data throughput to the host PC and introduce additional complexity in managing data flow within the FPGA. In future iterations, we may adopt an FPGA module with USB 3.0 communication capability and utilize onboard DDR3 memory for data buffering.

We also plan to replace the existing PC interface based on a Python script with a Graphical User Interface.

## 6. Conclusions

In this work, we present a novel open-source, compact, high-resolution biopotential amplifier designed for multiple applications and offered at a substantially lower cost than commercially available systems with comparable specifications. We demonstrate its performance in standard biopotential recordings as well as in specialized applications, including invasive electrophysiological measurements and fast neural electrical impedance tomography imaging.

This is particularly valuable for research laboratories and educational settings where high-resolution recording capabilities are required but available funding is insufficient to acquire commercial systems, or where detailed knowledge of the circuit architecture is essential for integration with other devices or modules.

## Data Availability Statement

All of the hardware and software is released under a GNU General Public License v3.0, with contribution and distribution welcomed. Electronic designs, firmware code and data acquisition scripts, are available on the UCL EIT group GitHub page: https://github.com/EIT-team/NeuroADC.

## Acknowledgements

This project and author E. Ravagli are supported by a UKRI Horizon MSCA Guarantee Postdoctoral Fellowship (UKRI Grant EP/X03691X/1). Author E. Ravagli thanks Dr. Dai Jiang for feedback on the design and preliminary iterations of the circuit board, and Dr. Florencia Maurinho-Alperovich for support in performing the animal experiment reported in this study.

